# Time-resolved single-cell RNAseq profiling identifies a novel *Fabp5*-expressing subpopulation of inflammatory myeloid cells in chronic spinal cord injury

**DOI:** 10.1101/2020.10.21.346635

**Authors:** Regan Hamel, Luca Peruzzotti-Jametti, Katherine Ridley, Veronica Testa, Bryan Yu, David Rowitch, John C. Marioni, Stefano Pluchino

## Abstract

Innate immune responses following spinal cord injury (SCI) participate in early secondary pathogenesis and wound healing events. Here, we used time-resolved scRNAseq to map transcriptional profiles of SC tissue-resident and infiltrating myeloid cells post-SCI.

Our work identifies a novel subpopulation of *Fabp5^+^* inflammatory myeloid cells, comprising both resident and infiltrating cells and displaying a delayed cytotoxic profile at the lesion epicentre, which may serve as a target for future therapeutics.

## Main Text

Traumatic spinal cord injury (SCI) is a debilitating central nervous system (CNS) pathology that lacks effective restorative treatments. The mechanisms of tissue recovery after SCI are dysregulated in comparison to normal wound healing, leading to a chronic wound state, rather than a return to homeostasis. While this state is characterized by persistent inflammation driven by CNS resident microglia (MG) and infiltrating myeloid cells^1^, the precise roles of myeloid cell subsets after SCI remains unclear. In particular, upon crossing the blood-brain barrier, tissue infiltrating monocyte-derived macrophages (MCd) take on the morphology of MG, and upregulate canonical MG markers^2^, making the two populations practically indistinguishable.

Here, we utilized a fate-mapping approach to investigate the ontology and temporal dynamics of spinal cord MG and infiltrating myeloid cells via FACS isolation and single-cell RNA sequencing (scRNAseq) in a mouse model of thoracic contusion SCI (Fig. 1A).

**Fig. 1.**
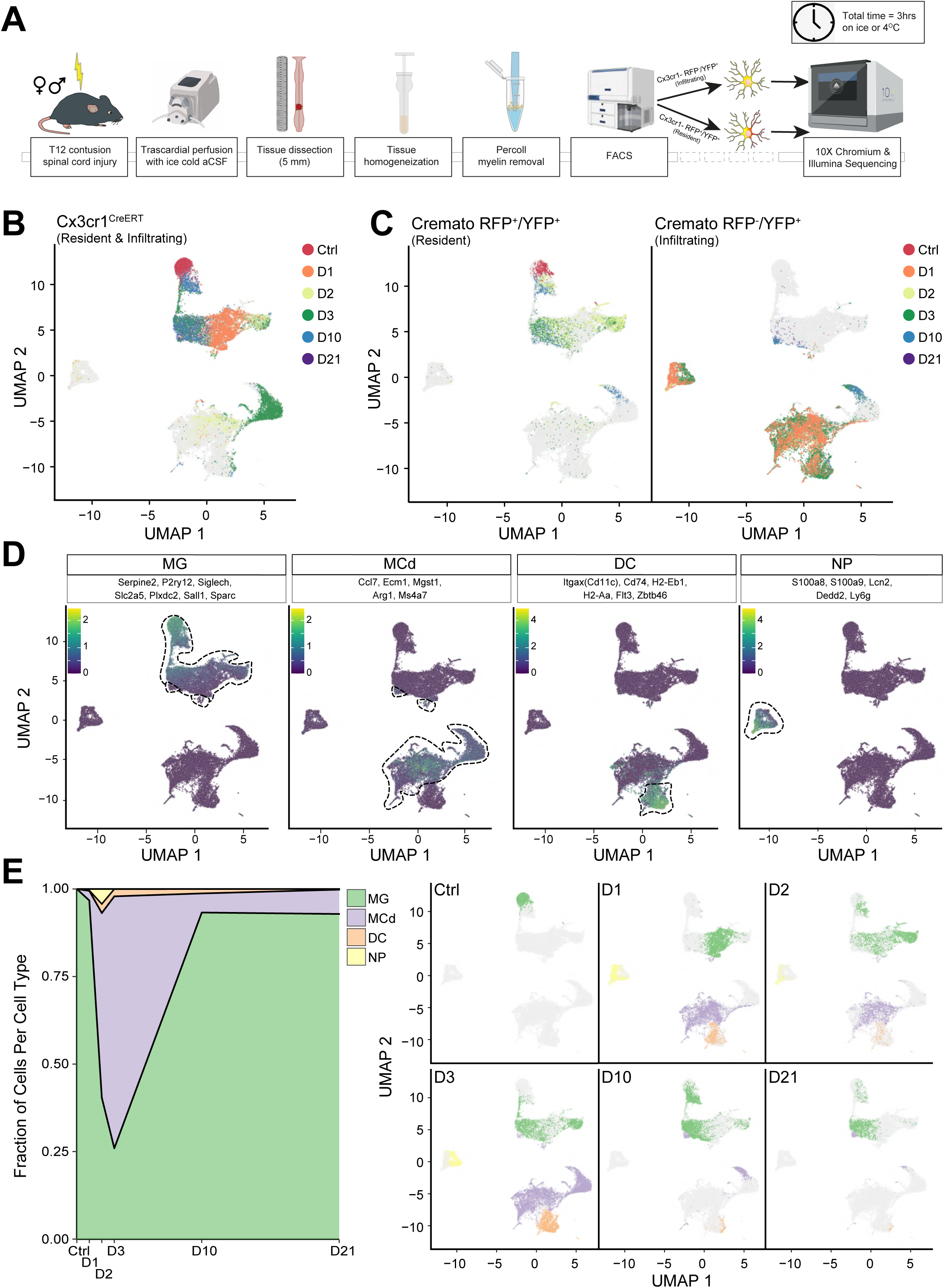
Characterization of CNS infiltrating and resident myeloid cells upon SCI. **A**, Myeloid lineage-specific cell isolation protocol used in this study. The illustration depicts the *Cremato* mouse line FACS only. **B**, UMAPs of the batch corrected dataset separated by strain and fate mapping. Each dot represents a cell and is coloured by the time of collection. **C**, UMAPs with cells (dots) coloured by the gene expression of the listed cell-type markers. Scale bars are the averaged log2-transformed counts from the listed marker genes. **D**, Area plot of the myeloid cell types comprising the *Cx3cr1* dataset (left) and UMAP of the entire dataset (right) separated by time point and coloured by cell type.

We first employed a G-protein coupled chemokine (C-X3-C motif) receptor 1 (Cx3cr1)^CreERT2^ mouse line, which expresses YFP under the endogenous Cx3cr1 promoter, to isolate and scRNAseq all YFP+ myeloid cells from the spinal cord of mice in the acute (day[D] 1, 2, 3), subacute (D10), and early chronic (D21) phases of SCI (Fig. 1B). Uninjured - laminectomised-only - mice were used as controls (Ctrl). We identified transcriptionally heterogeneous cell populations, which we hypothesized to include both CNS resident and infiltrating myeloid cells.

To directly identify the different myeloid cell subpopulations in the spinal cord, we adopted a Cx3cr1^CreERT2^:R26^tdTomato^ fate mapping mouse line (hereafter called *Cremato*), which exploits the differing lifespans of CNS resident *vs.* infiltrating myeloid cells^3, 4^. This allowed us to distinguish myeloid cells as double-positive (RFP+/YFP+) CNS resident and single-positive (RFP^−^/YFP^+^) CNS infiltrating (Fig. 1C), and to quantify their respective spatial distribution over time (**Extended Data Fig. 1**).

In total, we analysed 30,958 cells from Cx3cr1^CreERT^ and *Cremato* mice (**Extended Data Table 1**) and observed neither behavioural nor transcriptional differences between males and females post-SCI, allowing us to pool these datasets (**Extended Data Fig. 1**)^2, 5, 6^. Additionally, we observed lower-levels of dissociation- and FACS-induced gene expression^7^ in Ctrls, which suggests the protocol described here minimizes cellular activation and is optimal for experimental designs involving *ex vivo* cellular isolation and gene expression analyses (**Extended Data Fig. 1**)^6, 8, 9^.

After batch correction, we performed unsupervised clustering and visualised the output using Uniform Manifold Approximation and Projection (UMAP) (**Extended Data Fig. 1**). The cell types comprising each cluster were then determined based on fate mapping labels, Gene Ontology (GO), and core gene signatures from previous CNS scRNAseq studies (Fig. 1D)^2, 3, 9, 10^.

We confirmed the presence of previously described cell types and the myeloid-lineage specificity of the *Cremato* mouse line *in vivo* via single-molecule fluorescent in situ hybridization (smFISH) and confocal immunofluorescence microscopy (**Extended Data Fig. 2**). We found that Ctrls comprised MG only, as expected (Fig. 1E and **Extended Data Fig. 2**). Upon SCI, CNS infiltrating cell types, namely MCds, neutrophils (NPs)^11^, and myeloid dendritic cells (DCs), increased considerably in both the acute and subacute phases (Fig. 1E and **Extended Data Fig. 2**). In contrast, during the chronic phase (D21), the vast majority of isolated myeloid cells were either MG or MCds. We did not observe expression profiles characteristic of other CNS-associated cell types, indicating that our isolation methods were specific for our target population (**Extended Data Figs. 1** and **2**).

Recently, scRNAseq has been used to describe MG in development, homeostasis, and disease. A common feature across several CNS diseases is the downregulation of canonical MG markers upon activation (Fig. 2A)^2, 3, 5^. Indeed, we found that MG-specific markers *P2ry12*, *Plxdc2*, *Sall1*, *Siglech*, *Sparc*, and *Serpine2* were expressed in Ctrl MG but downregulated in MG by D1, as described^2, 3, 12^ (**Extended Data Fig. 3**). By D1, *Tmem119*, *Olfml3*, and *Grp34* were also downregulated in MG but upregulated in MCd (**Extended Data Fig. 3**). Other MG markers - *Fcrls*, *C1qa*, *Aif1/Iba-1*, *Hexb*, *Trem2* - were all stably expressed in MG, but increased in CNS infiltrating myeloid cells (**Extended Data Fig. 3**), which further supports previous findings of MCds adopting a MG-like phenotype after entering the CNS^2^. Interestingly, we found *S100a11* to be a novel marker for CNS infiltrating myeloid cells. *Chil3/Ym1* was expressed by MCds/NPs^13^, and *Ms4a7*^2^, *Ccl7*^14^, *Arg1*^3^, and, to a lesser extent *Mgst1*^10^, were restricted specifically to MCds (**Extended Data Fig. 3**). Using smFISH, we confirmed the dynamic post-SCI expression of *Serpine2* and *Ms4a7* by MG and CNS infiltrating myeloid cells, respectively (**Extended Data Fig. 4)**. Together these findings indicate that our myeloid cell scRNAseq dataset is consistent with and expands upon previous scRNAseq studies in other disease models.

**Fig. 2.**
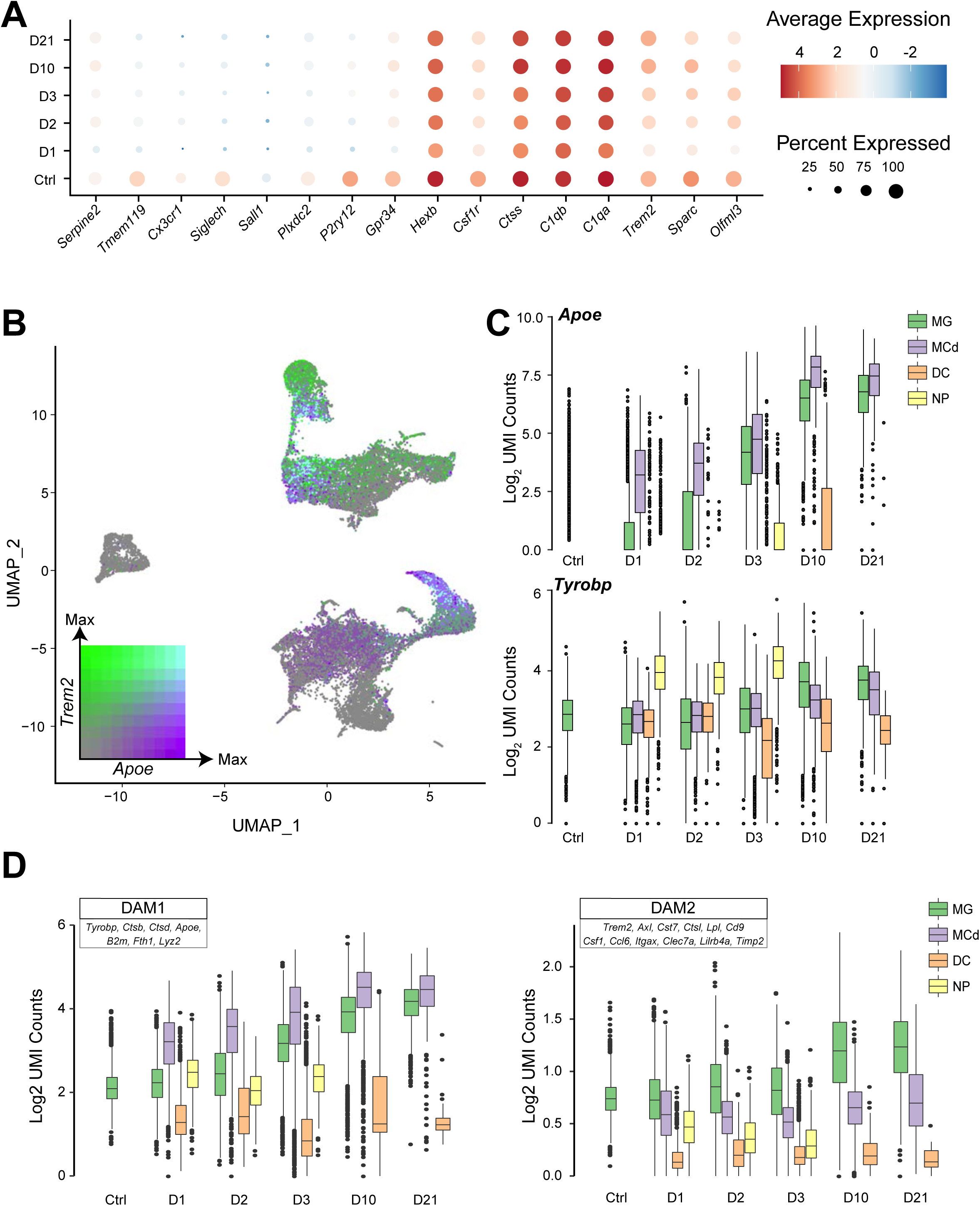
Validation against published myeloid cell studies. **A**, Seurat dot plot showing canonical MG marker expression. Scale bar is the natural log of the average log2-transformed counts. **B**, UMAP coloured by the scaled log2-transformed counts of *Apoe* and *Trem2*. **C**, Boxplots by time and cell type demonstrating the upregulation of *Apoe* (top) and *Tyrobp* (bottom) upon SCI. **D**, Boxplots by time point coloured by cell type, quantifying the average log2-transformed counts of DAM1- *vs*. DAM2-associated genes expressed per cell.

To gain further insight into the cellular and molecular mechanisms driving the observed transcriptional changes, we focused on the TREM2-induced APOE pathway, which has been shown to mediate the switch from homeostatic to activated myeloid cells^15^. Upon SCI, both MG and infiltrating myeloid cells upregulated apolipoprotein E (*Apoe*) and the TREM2-APOE pathway component, transmembrane immune signalling adaptor (*Tyrobp*) (Fig. 2B-C and **Extended Data Fig. 4**).

The upregulation of *Apoe* and *Tyrobp*, along with the downregulation of several homeostatic MG markers, has recently been described as the first step towards the acquisition of a neuroprotective, phagocytic, ‘*Disease Associated MG*’ (DAM) phenotype in murine models of Alzheimer’s disease and amyotrophic lateral sclerosis^12^. DAMs comprise two sequential stages (DAM1 and DAM2), which are conserved between mice and humans^12^, but their transcriptional profiles have yet to be described in SCI. In the acute phases post-SCI, we found that the expression of DAM1-associated genes increased in both MG and MCds. Conversely, the expression of DAM2-associated genes increased over time in MG only (Fig. 2D), with all but two DAM2-associated genes upregulated in MG by the subacute and early chronic phases post-SCI.

Next, we sought to identify new molecular processes involved in the immune response to SCI by using unsupervised analytical approaches. To this end, we first turned to unsupervised clustering results and generated cluster-specific differentially expressed genes followed by GO enrichment analysis. We noted that the fraction of myeloid cells per cluster changed over time and the correlation between the time of collection and the assigned cluster was significant (**Extended Data Fig. 5**), which inspired us to perform unsupervised trajectory analysis (Fig. 3A). Through our temporally specific samples (Fig. 3B), we gave directionality to this trajectory analysis and generated a map of MG and MCd transcriptional profiles across the secondary injury (**Extended Data Fig. 6**).

**Fig. 3.**
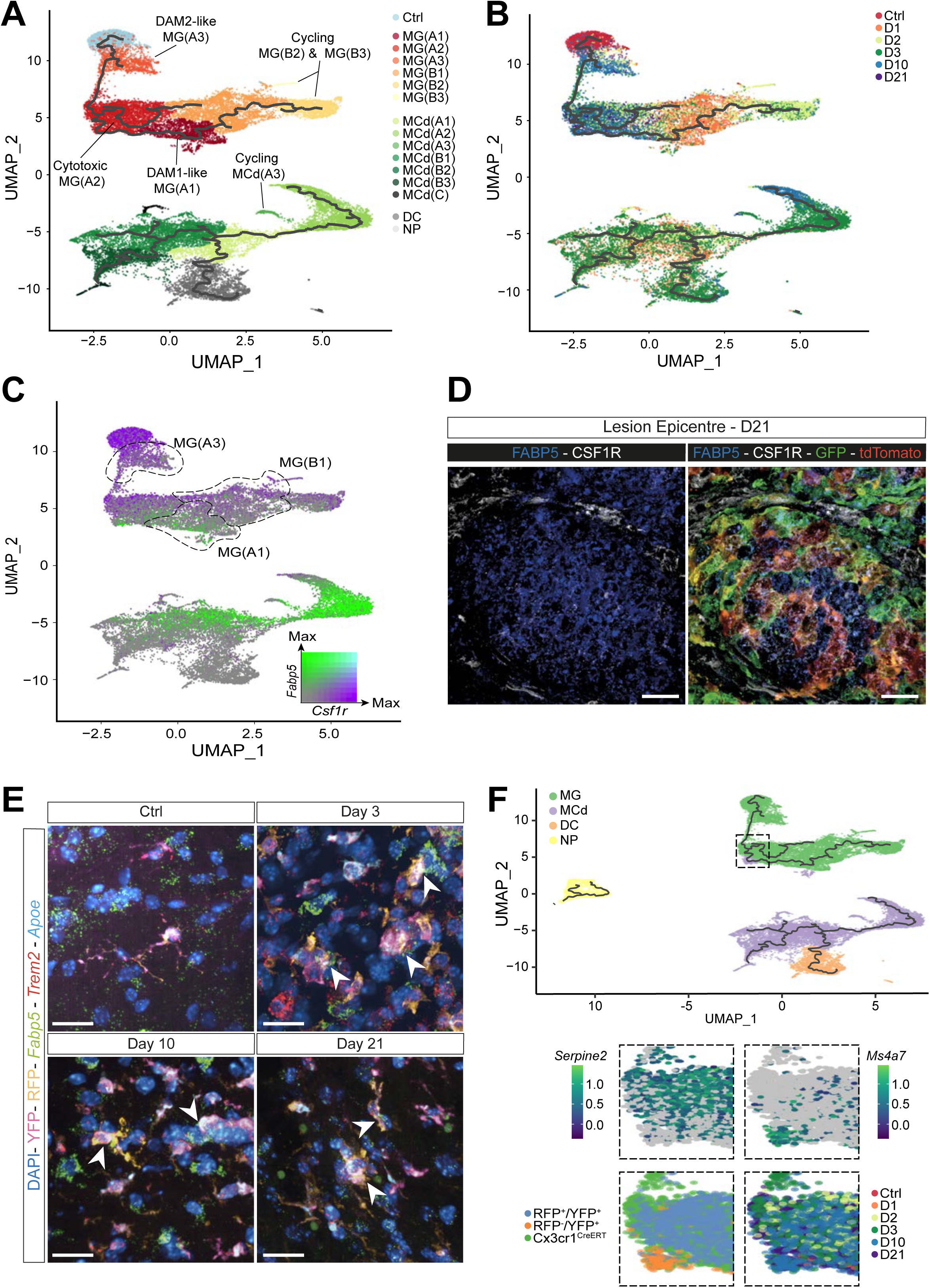
Dynamic trajectories of myeloid cells post-SCI. **A**, UMAP coloured by cluster, as determined by Leiden community detection using Monocle 3. Clusters are named based on their majority cell-type and their position in the trajectory. The UMAP is superimposed with the unsupervised reversed graph embedding-derived trajectory inferred via Monocle 3. **B**, UMAP coloured by time post-SCI, giving directionality to the trajectory. Trajectories in **A** and **B** are represented by the black lines. **C**, UMAP coloured by the scaled log_2_-transformed counts of *Fabp5* (green) or *Csf1r* (purple). Dotted lines depict relevant clusters. **D**, D21 expression of FABP5 and CSF1R at the lesion epicentre. Nuclei were stained with DAPI. Scale bars: 20 μm. Full panels in Extended Data 8. **E**, smFISH validation of the expression of *Fabp5, Apoe and Trem2*. Arrowheads indicated RFP^+^/YFP^+^ MG expressing *Fabp5*. Nuclei were stained with DAPI. Scale bars: 20 μm. Full panels in Extended Data 7. **F**, UMAP coloured by cell type with close up views of the subacute/early chronic MCd, RFP^−^/YFP^+^ cells which cluster with MG(A2) but retain the expression of the MCd marker, *Ms4a7*, and do not express *Serpine2*.

At D1, the majority of MG were in a highly activated MG(A1) or MG(B1) state characterized by GO terms for the production of (and response to) cytokines, recruitment of blood-borne leukocytes, phagocytosis, response to low-density lipoproteins, lipid transport and localization, and positive regulation of astrocyte activation, as described^1^ (**Extended Data Fig. 5**). However, MG(A1) and MG(B1) states differed in their expression of colony-stimulating factor 1 receptor (*Csf1r*), a gene required for MG proliferation and survival^16^, epidermal fatty acid-binding protein 5 (*Fabp5*), a target of the Apoe-induced transcription factor Bhlhe40*^15, 17^*, and DAM1 genes (**Extended Data Figs. 5** and **7**). Thus, MG(A1) were defined as DAM1-like, *Csf1r^lo^/Fabp5^+^/Trem2^−^*, while MG(B1) were *Csf1r^int^/Fabp5^lo^/Trem2^int^*. We observed the mutually exclusive expression of *Fabp5* and *Csf1r* in all myeloid cells and confirmed this observation at the protein level by confocal immunofluorescence microscopy at D21 (Fig. 3C-D and **Extended Data Figs. 5** and **8**). Furthermore, MG(B1) were found to express S-phase genes and GO terms suggesting a state of cell cycling initiation (**Extended Data Fig. 5**). Finally, compared to MG(B1), MG(A1) showed higher fold enrichments for GO-terms related to activation of the NF-κB pathway and lipid transport/localization, in line with previous findings on *Fabp5*^18^.

By D2-D3, MG(B1) progressed into one of three states: (i) the recently described cycling population^4^, MG(B2) and MG(B3), identified by *Trem2^int^* and *Mki67*, G2-M phase genes, and mitosis related GO-terms; (ii) MG(A1); or (iii) MG(A2) (***Extended Data Fig. 5***). Similarly, MG(A1) mainly transitioned into the MG(A2) state, through a *Fabp5hi/Trem2lo* trajectory (***Extended Data Fig. 5***). Notably, MG(A2), which were characterized as *Csf1r^int^/Fabp5^+^/Trem2^hi^/Apoe^hi^*, appear to be a novel cytotoxic, phagocytic, lipid-processing cell population with continued astrocyte activation, blood-borne leukocyte recruitment, and the production of interleukin-6, a cytokine known to sustain chronic inflammation^19^ (Fig. 3C-E; **Extended Data Figs. 5-8**. This cluster persisted from D2-D21 and while a small subset at D10 progressed into a *Csf1r^hi^/Fabp5^−^/Trem2^hi^* MG(A3) state, characterised by a neuroprotective DAM2 phenotype12 (**Extended Data Fig. 5**), most MG remained in a chronic, cytotoxic MG(A2) state. Quadruple immunofluorescence of FABP5, CSF1R, and *Cremato* fate-mapping in the lesion epicentre at D21 confirmed that CSF1R-/FABP5^+^ MG and MCds, likely corresponding to MG(A2), dominated the lesion core, while the majority of CSF1R^+^ MG, possibly MG(A3), remained perilesional (**Extended Data Fig. 8**). Furthermore, despite the D21 decrease in *Fabp5* observed via smFISH and scRNAseq, FABP5 expression was sustained at D21, suggesting it plays a persistent role in the cytotoxicity of the MG(A2) population post-SCI (Fig. 3C-E and **Extended Data Figs. 6-8**).

Finally, we focused on the dynamics of MCds and found that at D1, MCd occupy one of two states: MCd(A1) or MCd(B1). Both MCd clusters, based on our quantification and the presence of C-C Motif Chemokine Receptor 2 (*Ccr2*)^20^ (**Extended Data Figs. 1** and **5**), had yet to fully infiltrate the spinal cord parenchyma and differentiate into macrophages, as reported for this SCI stage^1^. MCd(A1) or MCd(B1) clusters showed enrichment for GO terms related to astrocyte activation, response to cytokines and reactive oxygen species (ROS), oxidative stress, respiratory bursts, phagocytosis, cellular detoxification, and lipid processing^14^. However, MCd(B1) showed GO terms for blebbing and seemed to be fated towards cytokine production and apoptosis as described^21^, while by D3 the fate of MCd(A1) correlated with its *Fabp5* expression (**Extended Data Fig. 5**). *Fabp5lo* MCd(A1) in fact differentiated into DCs, whereas *Fabp5hi* MCd(A1) transitioned towards either MCd(A2) or MCd(A3). MCd(A3) were a small cluster of proliferating MCds at D3 (**Extended Data Fig. 5**), as reported predominantly for MG, and to a lesser degree MCd, in SCI21. MCd(A2) was characterized as a cluster of infiltrated cells with GO terms for neurogenesis and wound healing but also cell killing, and both the positive and negative regulation of ROS and cytokine production. These opposing terms could be explained by the transition within MCd(A2) from a more pro-inflammatory *Fabp5hi* (D3) to a more neuroregenerative *Trem2^hi^* (D10) transcriptional profile (**Extended Data Fig. 5**), which we verified by smFISH (**Extended Data Fig. 7**). We observed a small group of RFP^−^/YFP+ MCd cells that clustered with *Fabp5+* MG(A2). We confirmed the presence of FABP5^+^/RFP^−^/YFP^+^ MCd, which accumulated at the lesion epicentre at D21 via confocal immunofluorescence microscopy (Fig. 3D and **Extended Data Fig. 8**). These D10/D21 cells remained *Serpine2^−^/Ms4a7^+^/Gpnmb^+^*, but otherwise adopted the transcriptional profiles of RFP^+^/YFP^+^ MG(A2) (Fig. 3F and **Extended Data Fig. 5**), suggesting that upon SCI, a subset of MCd infiltrate the CNS parenchyma and take on the transcriptional profiles, and potentially functions, of activated, cytotoxic MG.

Cumulatively, our data suggests that the myeloid cell response to SCI is highly dynamic with both beneficial and detrimental effects and inspired us to hypothesise that the downregulation of *Fabp5* expression would control the progression towards a homeostatic, potentially neuroprotective, DAM2-like phenotype.

Additionally, we identify a novel delayed cytotoxic transcriptional profile, *Fabp5^+^* MG(A2), which is adopted by both CNS resident and infiltrating myeloid cells and sustained at the lesion epicentre in the chronic phases of SCI. Whether or not this profile is transient, or can be targeted with specific therapies, requires further investigation.

Furthermore, our novel, publicly available scRNAseq dataset (https://marionilab.cruk.cam.ac.uk/SCI_Myeloid_Cell_Atlas/) complements and expands upon the findings of previous studies and will serve as a valuable resource for future investigations into the role of the immune responses after SCI.

## Acknowledgements

The authors wish to acknowledge Aviva Tolkovsky, Alice Braga, Jayden A. Smith, Yutong Chen, Tommaso Leonardi, Christian Frezza, Michele De Palma, and Giovanni Pluchino for their technical contributions and critical insights throughout the execution of the study. The authors acknowledge the Cambridge NIHR BRC Cell Phenotyping Hub for their cell sorting support and the CRUK Genomics Core Facility for processing the scRNAseq samples and for their technical support.

This work was supported by Wings for Life (RG 82921 to S.P.), the Bascule Charitable Trust (RG 75149 and RG98181 to S.P.), the Rosetrees Trust (A1850 to R.H. and S.P.), and by core funding from EMBL (to J.M.) and Cancer Research UK (C9545/A29580) to J.M), and the Wellcome Trust (D.H.R.). R.H was supported by the Cambridge Trust (10468562) and is the recipient of a Canadian Scholarship Trust Foundation, an MNI-Cambridge Douglas Avrith Graduate Studentship, and a Rosetrees Trust Studentship (A1850). L.P.-J. was supported by a senior research fellowship FISM - Fondazione Italiana Sclerosi Multipla - cod. 2017/B/5 and financed or co-financed with the ‘5 per mille’ public funding, and the Addenbrooke’s Charitable Trust (RG 97519).

## Author Contributions

R.H., L.P.-J. and S.P. conceived the study and designed the experiments; R.H. and J.M. designed the bioinformatics analyses and R.H. performed analyses under J.M. supervision. R.H. and B.Y. performed the animal experiments and optimized the cell isolation protocol. R.H., B.Y., and V.T. prepared the animal tissue for scRNAseq and tissue pathology. V.T. and R.H. performed the IHC experiments and analysis with the advice of L.P.-J. K.R. performed the smFISH experiments. R.H. performed the bioinformatics analysis and wrote the original manuscript, which was reviewed and edited by L.P.-J., J.M., and S.P. Funding for the project was acquired by R.H., S.P. and J.M. All authors approved of the final version of the manuscript.

### Competing Interests

The authors declare no competing interests.

## Methods

### Contusion Models of Spinal Cord Injury

Male and female mice aged 6 to 8 weeks were bred and housed in pathogen-free conditions and provided chow and water *ad libitum*. The following strains were used in this experiment: Cx3Cr1CreERT2 (purchased in 2018 from Jax, strain name: *B6.129P2(Cg)-Cx3cr1tm2.1(cre/ERT2)Litt/WganJ*) to isolate Cx3cr1-YFP^+^ myeloid-lineage cells; Cx3cr1CreERT2 crossed with TdTomato flox (*Cremato*) (purchased in 2018 from Jax, strain name: B6.Cg-Gt(ROSA)26Sortm9(CAG-tdTomato)Hze/J) to distinguish between YFP+ infiltrating myeloid cells and double-positive RFP/YFP CNS resident microglia (MG), as recently described^3^; C57BL/6 (purchased as needed from Charles River), hereon referred to as WT, for tissue pathology. We administered *Cremato* mice with daily intraperitoneal injections of 125 mg/kg of body weight of Tamoxifen (Sigma Life Science) dissolved in corn oil for 5 consecutive days to activate the Cre-recombinase, labelling all CX3CR1^+^ cells as double-positive for RFP and YFP. We induced SCI after a 28-day wash-out period, during which peripheral monocytes were replaced by single positive YFP bone marrow-derived monocytes.

After deeply anaesthetizing the animals with isoflurane (4% induction, 2% maintenance) in oxygen (1.5 l/min) we provided buprenorphine (Temgesic, RB Pharmaceuticals) (pre- and post-operatively), applied ocular ointment to prevent the eyes from drying, shaved the hair on the back of the mice, and swabbed it with a germicide prior to surgery. We placed the animals in prone position and under a surgical microscope, created a dorsal midline incision over the thoracic vertebrae with a sterile scalpel and separated the paravertebral muscles from using a spring scissor (Fine Science Tools). We identified T12 as the apex of the dorsal aspect and performed the laminectomy at T12 using Dumont #2 laminectomy forceps (Fine Science Tools) and spring forceps (Fine Science Tools) leaving the dura intact. In order to maintain vertebral column stability, we did not remove the lateral part of the vertebra at the site of laminectomy. The extension of the laminectomy was consistent between animals at approximately 3 mm in width and 5 mm in length, sufficient to allow room for the 1.3 mm diameter impactor tip.

For the control group (Ctrl), we performed a laminectomy at T12 and omitted the contusion injury step. For the spinal cord injury (SCI) group, we induced a bilateral contusion injury on the exposed spinal cord at T12 using the Infinite Horizon (IH) impactor device (Precision Systems and Instrumentation, Lexington, KY) as previously described^22^. First, we aligned the mouse-impacting tip over the exposed cord and centred it over the central vein, avoiding any overlap with the transverse processes. Then a moderate contusion injury (70 kilodyne force) was induced using the IH impactor. After the injury, we closed the incision with 7-mm AutoClips (Fine Science Tool), which were removed after 14 days post-SCI. We expelled the bladders of mice subjected to SCI by applying manual abdominal pressure twice a day until they regained urinary function or reached their endpoint.

We performed all experimental animal procedures in accordance with the Animals (Scientific Procedures) Act 1986 Amendment Regulations 2012 following ethical review by the University of Cambridge Animal Welfare and Ethical Review Body (AWERB). Animal work was covered by the PPL 7008840 (to Stefano Pluchino).

### Motor Behavioural Assessment

We assessed hindlimb motor performance using the open-field Basso Mouse Scale (BMS)^23^, with scores ranging from 0 (complete hindlimb paralysis) to 9 (healthy). We scored the animals for 4 min in an open field prior to the SCI induction and 1, 3, 5, 7, 10, 14, 18, and 21-days post to ensure that the mice displayed the level of hind-limb locomotor impairment expected from a moderate contusion SCI. Only animals with left and right hindlimb BMS scores within 2 points of each other and a BMS score of 0 at day 1 post-SCI were used in this study. For statistical analysis of the BMS scores, we took the average scores of the left and right hind limbs resulting in a single BMS score for each animal.

### Spinal Cord Extraction for Tissue Pathology

Mice were deeply anaesthetized with intraperitoneal injections of 100 μl of pentobarbital sodium and transcardially perfused with ice-cold artificial cerebral spinal fluid (aCSF) for 7 minutes followed by ice-cold 4% paraformaldehyde (PFA) in PBS with a pH 7.4. We post-fixed the spinal cords in 4% PFA for 12 hours at 4°C and then washed with 1X PBS. After dissecting out the spinal cords, we cryoprotected them for at least 24 hours in 30% sucrose (Sigma) in 1X PBS at 4°C before embedding them in Optimal Cutting Temperature (OCT) Compound (VWR Chemicals) and snap freezing them with isopentane on dry ice. We stored frozen cord blocks at −80°C until a total of 15 mm of each spinal cord segment, centred on the lesion, was sectioned coronally at 20 μm thickness (cryostat CM1850; Leica) and collected onto SuperfrostPlus slides (ThermoScientific). We stored sections at −80°C until needed.

### Immunofluorescence

In preparation for immunohistochemistry, we baked spinal cord sections mounted on slides at 37°C for 30 min then washed with MilliQ water and PBS, for a total of 15 minutes. Then, we incubated the sections in blocking solution (1X PBS with 0.1% Triton X-100 (Sigma) and 10% normal goat serum (NGS; Sigma)) for 1 hour at room temperature (RT). We then incubated the sections overnight with the primary antibody in a solution of PBS 1X + 0.1% Triton X100 (Sigma) + 1% NGS at 4°C. Then, after 3 washes with PBS 1X + 0.1% Triton X100 for a total of 15 minutes, we incubated the slides for 1 hour in a humid chamber with the species-appropriate fluorochrome-conjugated secondary antibodies (1:1000 dilution) in a solution of PBS 1X + 0.1% Triton X100 + 1% NGS. We blocked the staining by washing the slides first 3 times with PBS 1X + 0.1% Triton X100 and then with PBS for a total of 20 minutes. We counterstained the sections with DAPI (Roche) diluted 1:10000 in PBS for 10 minutes at RT. Finally, we washed the sections with PBS and mounted them on glass coverslips with fluorescence mounting medium (Dako). We acquired images of the stained spinal cord sections using a vertical epifluorescence microscope (Leica DM6000) or a confocal microscope (Leica TCS SP5).

### Quantification

We acquired images of the stained spinal cord sections using a vertical epifluorescence microscope (Leica DM6000) at a magnification of 20X and spaced approximately 400 μm apart. We converted single 20X images to mosaics using ImageJ 1.53c and quantified cells using 2 ROIs of 700 μm^2^ per tissue section. We then quantified RFP and YFP positive cells by manual counting, while DAPI positive cells were quantified using CellProfiler. Quantification results were presented as a percentage of positive cells out of all DAPI positive cells. We used a total of n= 2 slices for each distance from the lesion epicentre.

### smFISH

Spinal cord sections were baked at 65°c for 1 hour and fixed in 4% paraformaldehyde (PFA) solution at 4°c for 15mins. Slides were then washed and dehydrated in PBS (1x) and ethanol gradients from 50% to 100% for a total of 30 mins. Slides were air-dried before automated single-molecule fluorescent in situ hybridization (smFISH) protocol. Multiplex smFISH was performed on a Leica BondRX fully automated stainer, using RNAScope Multiplex Fluorescent V2 technology (Advanced Cell Diagnostics, 322000). Slides underwent heat-induced epitope retrieval with Epitope Retrieval Solution 2 (pH 9.0, Leica AR9640) at 95°C for 5 mins. Slides were then incubated in RNAScope Protease III reagent (ACD 322340) at 42°C for 15 mins, before being treated with RNAScope Hydrogen Peroxide (ACD 322330) for 10 mins at RT to inactivate endogenous peroxidases. All double-Z mRNA probes were designed against mouse genes by ACD for RNAScope on Leica Automated Systems. Slides were incubated in RNAScope 2.5 LS probes for 2 hours at RT DNA amplification trees were built through consecutive incubations in AMP1 (preamplifier, ACD 323101), AMP2 (background reduction, ACD 323102) and AMP3 (amplifier, ACD 323103) reagents for 15 to 30 mins each at 42°C. Slides were washed in LS Rinse buffer (ACD 320058) between incubations. After amplification, probe channels were detected sequentially via HRP–TSA labelling. To develop the C1–C3 probe signals, samples were incubated in channel-specific horseradish peroxidase (HRP) reagents for 30 mins, TSA fluorophores for 30 min and HRP-blocking reagent for 15 min at 42 °C. The probes in C1, C2 and C3 channels were labelled using Opal 520 (Akoya FP1487001KT), and Opal 650 (Akoya FP1496001KT) fluorophores (diluted 1:500). The C4 probe complexes were first incubated with TSA–Biotin (Akoya NEL700A001KT, 1:250) for 30 min at RT, followed by streptavidin-conjugated Atto425 (Sigma 56759, 1:400) for 30 min at RT. Directly following the smFISH assay, tissue was incubated with anti-RFP (ThermoFisher, MA515257, 1:200) and anti-GFP (Abcam, ab13970, 1:400) antibodies in blocking solution for 1 hour. To develop the GFP antibody signal, samples were incubated in donkey anti-goat HRP (ThermoFisher - A15999, 1:200) for 1 hour, TSA biotin (Perkin Elmer - NEL700A001KT, 1:200) for 10 min and streptavidin-conjugated Alexa 700-streptavidin (Sigma - S21383, 1:200) for 30 min. To develop the RFP antibody signal, samples were then incubated in donkey anti-mouse Alexa 594 (ThermoFisher, A21203, 1:500). Samples were then incubated in DAPI (Sigma, 0.25 μg /ml) for 20 mins at RT, to mark cell nuclei. Slides were briefly air-dried and manually mounted using ~90 μl of Prolong Diamond Antifade (Fisher Scientific) and standard coverslips (24 × 50 mm^2^; Fisher Scientific). Slides were dried at RT for 24 hrs before storage at 4°C for > 24 hrs before imaging. SmFISH stained mouse spinal cord slides were imaged on an Operetta CLS high-content screening microscope (Perkin Elmer). Image acquisition was controlled using Perkin Elmer’s Harmony software. High-resolution smFISH images were acquired in confocal mode using an sCMOS camera and x40 NA 1.1 automated water-dispensing objective. Each field and channel was imaged with a z-stack of 20 planes with a 1 μm step size between planes. All appropriate fields of the tissue section were manually selected and imaged with an 8% overlap. To segment single cells and quantify RNA spots from high-resolution images, analysis scripts were created on Harmony software (Perkin Elmer) using customizable, predefined, function blocks. Maximum intensity z-projection images were generated from confocal z-stacks per field. Single nuclei were selected from DAPI signal, and doublet and partial nuclei discarded via a DAPI morphology-driven linear classification model. The cytoplasm of single cells was identified by intensity thresholding of RFP/YFP immunofluorescence signals in the surrounding region to single nuclei. RNA spots were automatically quantified across single cells. All cells with a spot count above five were considered positive for that gene.

### Spinal Cord Extraction and Cell Isolation for scRNAseq

Mice were deeply anaesthetized with intraperitoneal injections of 100 μl of pentobarbital sodium and transcardially perfused with ice-cold artificial cerebral spinal fluid (aCSF) for 7 min. After perfusion, we voided the spinal columns using a 5 ml syringe filled with ice-cold aCSF. We placed the spinal cords on a petri dish above millimetre graph paper and cut a 5 mm section centred on the lesion using a sterile blade. For the first 2 batches, mice were not pooled before sequencing. After establishing that sex- and condition-matched littermates showed no transcriptional differences, up to 3 mice were pooled in order to maximize cell yields for sequencing. We mechanically dissociated the tissue in a glass homogenizer with 6 ml of aCSF-based Homogenization buffer [aCSF plus 10 mM HEPES (Sigma) to maintain stable pH despite CO2 released from the cells, 1% BSA (Sigma) to reduce cell clumping, 1 mM EDTA (Thermo Scientific) to chelate Ca^++^ and Mg^++^, reducing cell adhesion, 10 mg/ml of DNAse (Roche) 3000U to degrade any free-floating DNA and reduce clumping of cells, and 40 units/ μl of RNAse inhibitor (Invitrogen) to minimize the degradation of the cells mRNA transcripts]. We chose mechanical dissociation over enzymatic methods because it is more rapid, can be performed at ice-cold temperatures, and is less likely to cause conformational changes of surface receptors, which could alter cell function.

After tissue dissociation, we filtered the suspension with a pre-wet 40 μm strainer and rinsed the homogenizer with 2 ml of Homogenization buffer to increase cell yield. To remove myelin and debris from the cell suspension, we added 2.7 ml of isotonic 9:1 Percoll (Sigma) to 10X PBS to each sample. We gently mixed the samples and centrifuged at 800 g for 20 minutes at 4°C with a brake speed of 0. Myelin debris visibly layered at the surface and was carefully removed with a pipette. To remove the remaining Percoll, we added an ice-cold buffer of 5% autoMACS Rinsing Solution (Miltenyi Biotec) in 1X MACS BSA solution (Miltenyi Biotec) and centrifuged at 800 g for 5 minutes at 4°C. Then we resuspended the pelleted cells in 200μl Fluorescent Activated Cell Sorter (FACS) buffer [see Homogenization Buffer ingredients but dissolved in Cell Staining Buffer (Biolegend) not aCSF, and with 7-AAD live/dead stain (Invitrogen) at a concentration of 1:50].

The *ex vivo* FACS samples were sorted using a BD FACS Aria III cell sorter set to 3-way purity with a 100 μm nozzle at 20 psi. As our panel consisted of RFP, YFP and 7-AAD, fluorochrome compensation was not required. All gates were set based on the following: unstained WT sample, 7-AAD stained WT sample treated with DMSO to kill the cells, and for the *Cremato* samples, non-tamoxifen treated *Cremato* mice (RFP^−^, YFP+) to control for bleed-through of the non-tamoxifen activated RFP.

### Single-cell isolation and sequencing

FACs-isolated cells were sequenced on a single-cell level using the microdroplet based protocol, 10X Genomics Chromium Single Cell 3’ Solution (v2 chemistry for D2 Cx3cr1 samples, v3 for all others). As per the manufacturer instructions, approximately 18,000 cells per sample were loaded into each well. Single cells were isolated with a gel bead in emulsion, and mRNA transcripts were barcoded by cell and transcript (via unique molecular identifiers) before the generation of barcoded libraries, which were sequenced with the NovaSeq 6000 (Illumina) to an average depth of at least 40,000 reads per cell and > 85% sequencing saturation, as calculated by Cell Ranger. Raw and processed data files are available on the GEO (https://www.ncbi.nlm.nih.gov/geo/query/acc.cgi?acc=GSE159638).

### Single Cell RNA-seq Data Pre-processing

Sequenced samples were demultiplexed and aligned to the GRCm38 (mm10) using the Cell Ranger 3.1.0 pipeline (10X Genomics). Notably, the gene Clec7a was not present on the mm10 dataset supplied by Cell Ranger, so we generated a custom Cell Ranger mm10 dataset to include this gene. We performed all scRNAseq data processing (post-Cell Ranger) in R version 3.6.3. All packages used are free of charge and publicly available at the Comprehensive R Archive Network (https://cran.r-project.org), the Bioconductor project (http://bioconductor.org), or their respective GitHub pages. Our complete R analysis workflow, including scripts for generating the figures, is available on https://github.com/regan-hamel/SCI_2020: regan-hamel/SCI_2020.

### Quality Control, Batch Correction & Clustering

This section is based on a combination of packages and workflows developed by Marioni group, with the exception of SoupX.

We first corrected each batch of unfiltered gene/cell matrices for the effects of barcode swapping (DropletUtils package) and then filtered out empty droplets (DropletUtils). As our scRNAseq samples originated from homogenized tissue, we anticipated high levels of free-floating, background mRNA levels even with barcodes corresponding to intact cells. We estimated and subtracted this background (SoupX). Then, we calculated per cell quality metrics (scater) and identified and removed low-quality cells using a first pass threshold of 600 UMI counts and 600 unique genes per cell.

We evaluated the need for batch correction by first comparing two condition-, litter-, sex-matched samples that were sequenced on the same day. We observed no differences in UMAP substructure (based on visual inspection) between samples. Next, we investigated two condition- and sex-matched samples from different litters, collected on different dates and observed shifts between UMAP substructures. Thus, we applied the mutual nearest neighbour batch correction approach between samples collected on different dates only. Before batch correction, we employed the package to perform multi-batch normalization (batchelor), modelled the variance (scran) and selected the highly variable genes to be used in the batch correction (batchelor). Batches with the greatest heterogeneity (i.e. the greatest mixture of condition and sex) were used to set the reference space for all the batches. We used the batch corrected data to visualize the entire dataset using Uniform Manifold Approximation and Projection (UMAP) (scater) with the default parameters. We clustered the dataset by building a shared nearest neighbour graph (scater) and used the walktrap algorithm (igraph) to identify clusters. Note these clusters were used for quality control purpose and were not used downstream. Clusters with a high percentage (> 10%) of total UMI counts corresponding to mitochondrial genes were removed; the average UMI counts in these clusters were generally lower than other clusters, suggesting these clusters corresponded to damaged and dying cells, rather than a population of cells with endogenous mitochondrial gene upregulation. We also removed clusters that both expressed canonical marker genes of more than one cell type and had a higher than average UMI counts, as they likely represented doublets. After removing low-quality clusters of cells, we re-performed the batch correction and clustering and used this post-quality control dataset for all downstream analysis.

### Trajectory Analysis

We performed trajectory analysis using the beta version of Monocle3. To prepare for the trajectory analysis, we processed the post-quality control dataset following the Monocle3 workflow. First, we converted the non-log transformed counts matrix to a cell dataset object (cds). Then we called preprocess_cds with the default parameters to normalize the dataset and perform principal component analysis. We re-performed the mutual nearest neighbour batch correction (batchelor) on this cds by calling align_cds and correcting, as before, only between samples collected on different dates. Next we computed the UMAP with the following parameters: minimum distance = 0.1; metric = cosine; number of neighbours = 15. We performed unsupervised clustering via Leiden community detection by calling cluster_cells on the batch corrected cells with the parameters reduction_method = UMAP and the number of nearest neighbours (k)=16. Finally, we built the trajectory by calling the function learn_graph with the parameters use_partitions = TRUE and close_loop = FALSE. Note that for the downstream analysis we used only the cluster membership, trajectory, and UMAP, but not the normalized gene expression matrix from this Monocle3-processed dataset.

## Differential Gene Expression & Gene Ontology Enrichment Analysis

We generated differentially expressed genes (DEGs) for each cluster by calling the scran function, findMarkers, on the multi-batch normalized log2-transformed counts (batchelor; as described in the *Quality Control, Batch Correction & Clustering* section). The findMarkers function identifies DEGs by pairwise Welch t-Tests comparisons corrected for multiple testing via the Benjamini-Hochberg method and combines the comparisons into a ranked list of markers for each cluster. We used these DEGs for Gene Ontology (GO) enrichment analysis. To this end, we input the significant DEGs (FDR< 0.0005, FC> 1.25, ranging from 455-848 genes) for each cluster into the PANTHER Classification System overrepresentation test (release 2020-06-01) following the instructions on the Gene Enrichment analysis page (http://geneontology.org/docs/go-enrichment-analysis/), including the optional use of a custom reference list. P-values computed using the default setting of Fisher’s Exact Test were corrected for multiple testing using the Benjamini-Hochberg method. We examined only biological processes that were over-represented.

### Cell Type Annotation

We assigned a cell type identity to each cluster using its top DEGs, GO terms, and the expression of canonical marker genes. We then refined these annotations, for example in the case of MG(A2), which contained both MG and MCds, using the CNS resident *vs.* CNS infiltrating labels from our fate-mapping *Cremato* mouse line.

### Dissociation and FACS-associated Expression Analysis

For Extended Data Figure 1C, we analysed the expression of dissociation and FACS-associated gene expression across several studies. The genes considered were those identified by van den Birk et al., 2017^7^ that intersected with the genes of all 5 datasets: *Mt1, Ier2, Dnaja1, Socs3, Atf3, Jund, Ppp1r15a, Hspe1, Dusp1, Nfkbia, Hsph1, Jun, Junb, Egr1, Ubc, Zfp36, Hsp90aa1, Mt2, Dnajb1, Btg2, Nr4a1*. We downloaded gene-matrix text files from Genome Expression Omnibus, read them into R and converted them to Single Cell Experiment objects (SingleCellExperiment). We removed cells which did not meet the following quality control (including in our data): greater than 600 UMI counts, greater than 600 unique genes per cell, and less than 10% counts from mitochondrial genes. Notably, samples differed in their platforms and disease states, but all cells analysed were MG. The following samples were used: Yang 2018^24^ (Drop-seq, whole-brain enzymatic dissociation, C1q^+^ Sham control MG), Linnarsson 2018^6^ (10X Chromium, whole-brain mechanical homogenization, homeostatic MG), Stevens 2019^5^ (10X Chromium, mechanical homogenization, FACS of homeostatic MG – note this dataset consisted only of highly variable genes and all mitochondrial genes had been removed), Movahedi 2019^9^ (10X Chromium, whole-brain enzymatic dissociation, homeostatic MG), and Ctrl RFP^+^/YFP^+^ MG from the current study.

### Statistics

We analysed the statistical significance of the BMS scores between males and females using an unpaired two-tailed t-Test. We performed statistical analyses on the scRNAseq data in R. We analysed the significance of the downregulation of MG-specific genes (S. Fig 2) using a Welch t-Test corrected for multiple testing via the Benjamini-Hochberg method based on the total number of genes in the dataset (scran). We considered differences to be significant with the P values were < 0.05. To test for independence between the categorical variables, cluster membership and time point, we used Pearson’s chi-squared test for independence. First, we generated a contingency table of cluster vs time point, then tested for independence using Pearson’s chi-squared test by calling chisq.test (stats) on the table. To understand how each comparison contributed to the significant result, we investigated the standardized residuals. To determine if a standardized residual was significant, we squared them to obtain chi-squared values then called pchisq (stats) with df= 1 and lower.tail= FALSE to obtain a P-value, which we compared with the Bonferroni corrected P values as described^25^.

## Notes

https://www.ncbi.nlm.nih.gov/geo/query/acc.cgi?acc=GSE159638

https://marionilab.cruk.cam.ac.uk/SCI_Myeloid_Cell_Atlas/

https://github.com/regan-hamel/SCI_2020

### Competing Interest Statement

The authors have declared no competing interest.

