## Extended Data Figures 1-4 for "Time-resolved single-cell RNAseq profiling identifies a novel *Fabp5*-expressing subpopulation of inflammatory myeloid cells in chronic spinal cord injury"

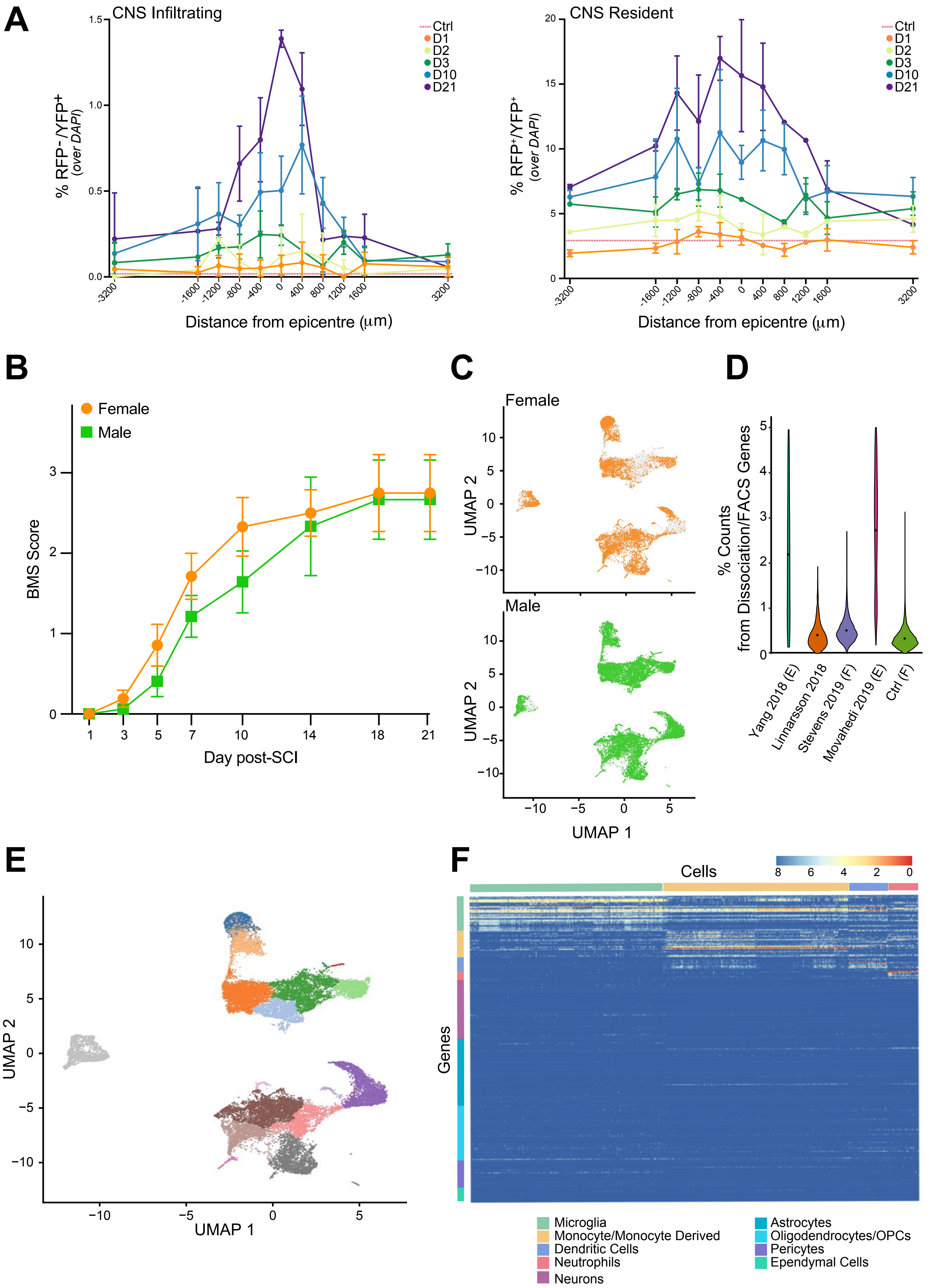


Hamel et al., Extended Data Figure 1

**Extended Data Fig. 1. Quality controls. A,** Quantification of $RFP^-/YFP^+$ infiltrating myeloid cells (left) or $RFP^+/YFP^+$ CNS resident myeloid cells (right) as a proportion of $DAPI^+$ at set distances rostral (+) or caudal (-) of the lesion epicentre. Data are mean % (± SD) and have been collected from n= 3 mice per time point. **B,** Basso Mouse Score (BMS) from male and female mice. Data are mean numbers (± SEM) and have been collected from n ≥ 4 mice per time point. **C,** UMAPs of cells from male *vs*. female mice. **D,** Violin plots showing the percentage of all UMI counts originating from dissociation and FACS associated genes, per Ctrl MG cell, across several droplet-based scRNAseq studies after applying the same quality control metrics. Black dots represent the mean. F= FACS, E= enzymatic dissociation (*vs.* mechanical). **E,** UMAP of the entire dataset, coloured by cluster, as determined by Leiden community detection using Monocle 3. **F,** Heatmap of the $log_2$-transformed counts of canonical marker genes for CNS-associated cell types.

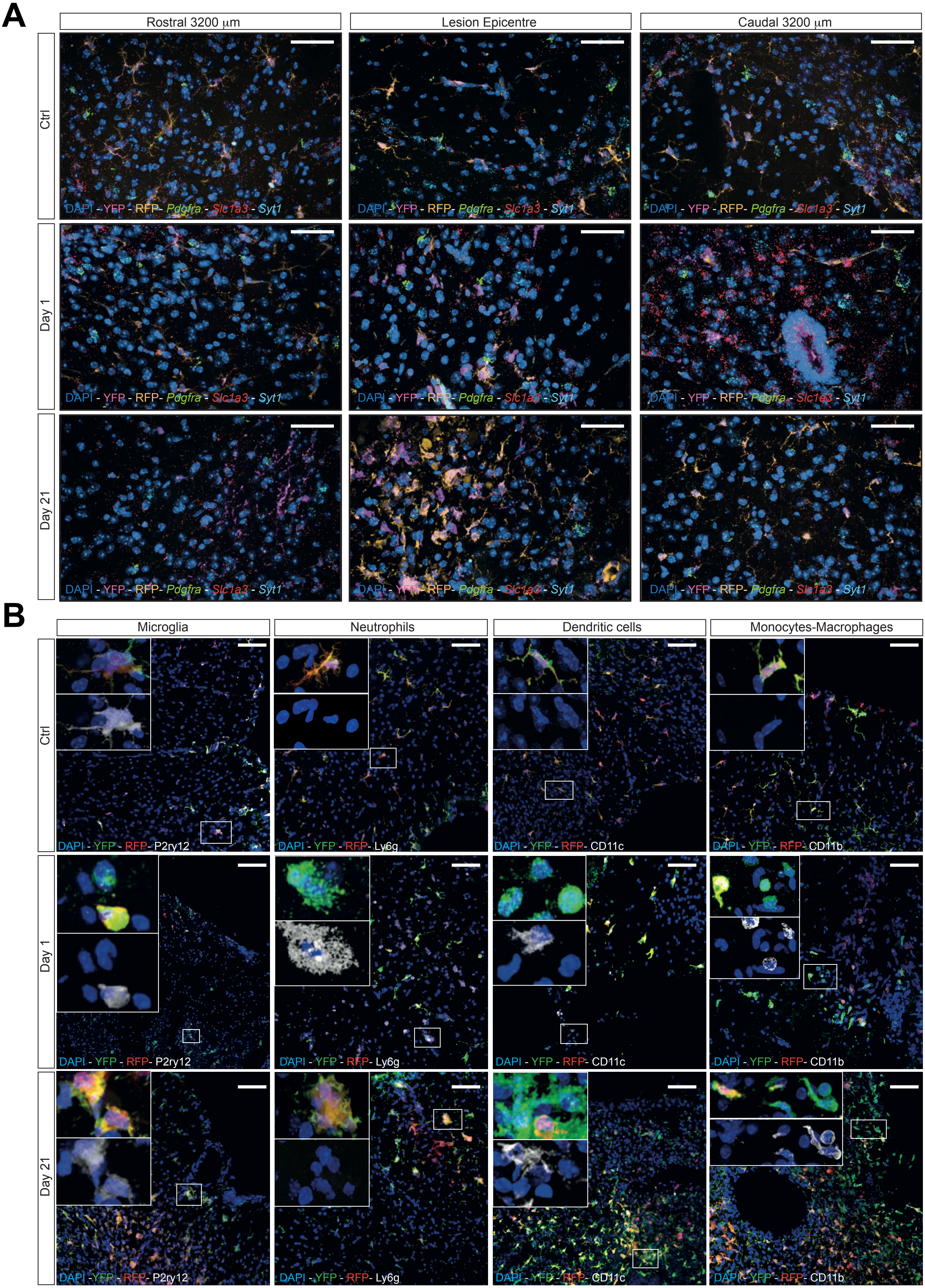


Hamel et al., Extended Data Figure 2

**Extended Data Fig. 2. SmFISH and confocal immunofluorescence microscopy validation of cell type identification data in Fig. 1. A,** Expression of cell-type markers *Pdgfra* (oligodendrocyte progenitor cells), *Slc1a3* (astrocytes), and *Syt1* (neurons) from Ctrl, D1, and D21. **B,** Expression of the commonly accepted markers P2ry12, Ly6g, CD11c and CD11b by microglia, neutrophils, dendritic cells and monocytes-macrophages, respectively, at the lesion epicentre. Nuclei were stained with DAPI. Scale bars: 60 μm.

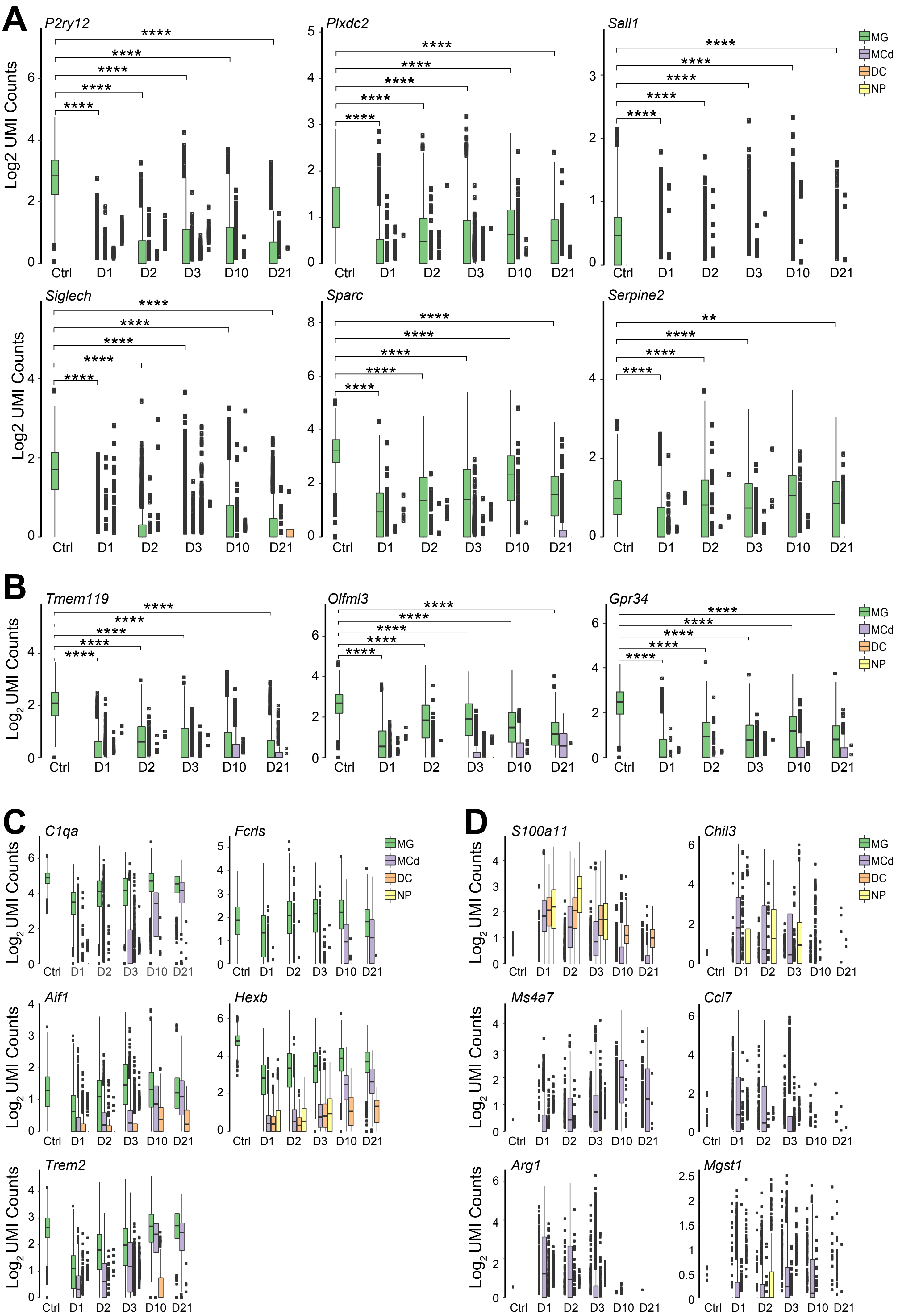


Hamel et al., Extended Data Figure 3

**Extended Data Fig. 3. Temporal dynamics of select gene expression by myeloid cells upon SCI. A,** Boxplots by cell type and time point showing the $log_2$-transformed expression of MG marker genes that remain exclusive to MG. Welch t-test with Benjamini-Hochberg post-hoc correction; ns $p \geq 0.05$; *$p < 0.05$, **$p \leq 0.01$, ***$p \leq 0.001$, ****$p \leq 0.0001$). **B,** Boxplots by cell type and time point demonstrating the upregulation of select MG makers in MCd upon SCI. Statistical analysis as described in **A**. **C,** Boxplots by time and cell type of several MG markers, including *Trem2*, which were upregulated in various myeloid cell types upon SCI. **D,** Boxplots by cell type and time point of genes that distinguish CNS infiltrating myeloid cells from resident MG. Bins in each boxplot are ordered from left to right: MG, MCd, DC, NP.

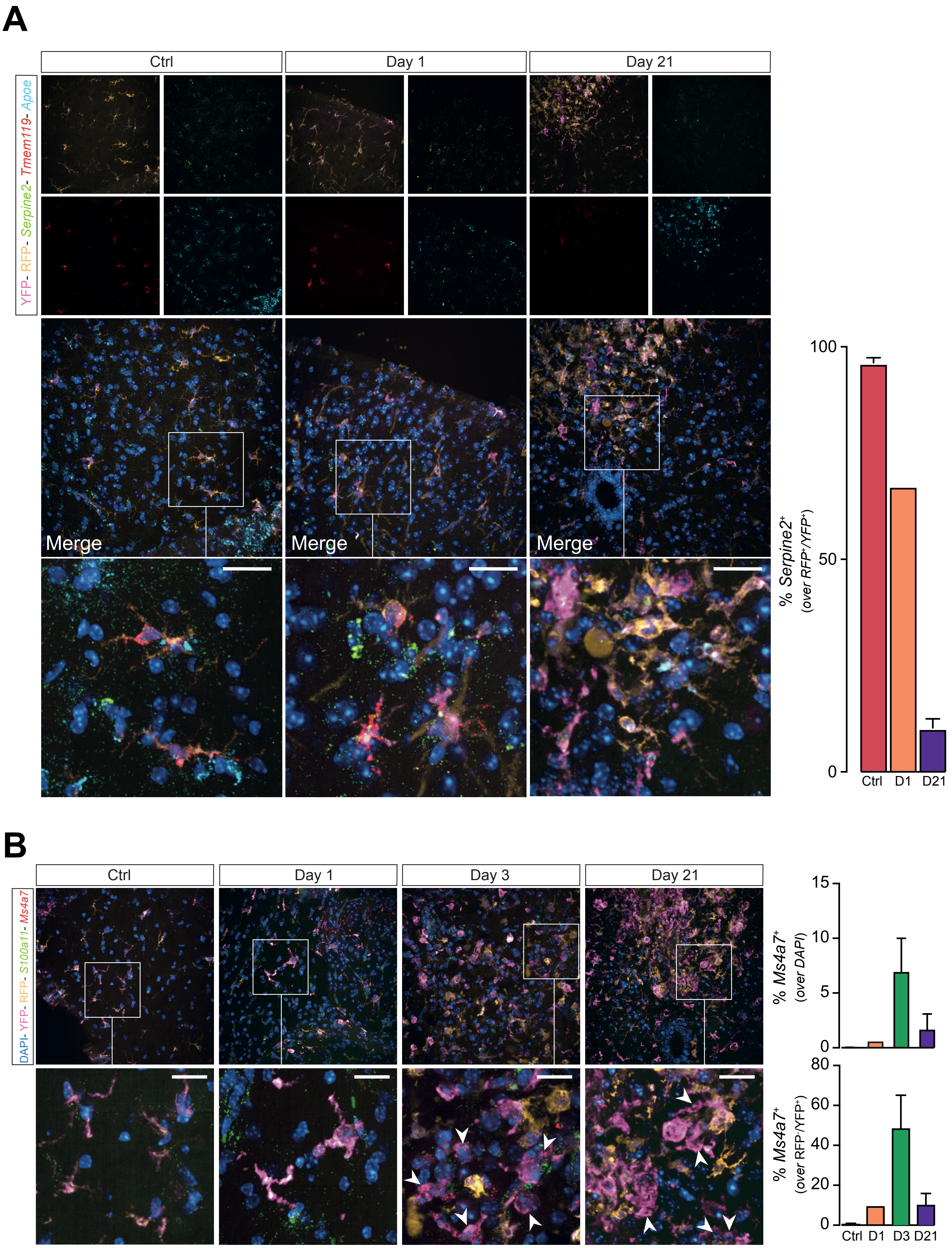


Hamel et al., Extended Data Figure 4

**Extended Data Fig. 4. SmFISH validation of select data from Extended Data Fig. 3**. **A,** Expression of *Tmem119*, *Serpine2* and *Apoe* from Ctrl, D1, and D21. The bar graph shows the quantification of *Serpine2*$^+$ cells out of all RFP$^+$/YFP$^+$ cells. **B,** Expression of *Ms4a7* and *S100a11* from Ctrl, D1, D3 and D21. The bar graphs show the quantification of *Ms4a7*$^+$ cells out of either all cells (top graph) or all RFP$^-$/YFP$^+$ cells infiltrating myeloid cells (bottom graph). Arrowheads indicate *Ms4a7*$^+$/RFP$^-$/YFP$^+$ cells. Data are mean % (± SD) from n= 2 mice per time point. Nuclei were stained with DAPI. Scale bars: 20 μm.
