## Extended Data Figures 5-8 + Table 1 for "Time-resolved single-cell RNAseq profiling identifies a novel *Fabp5*-expressing subpopulation of inflammatory myeloid cells in chronic spinal cord injury"

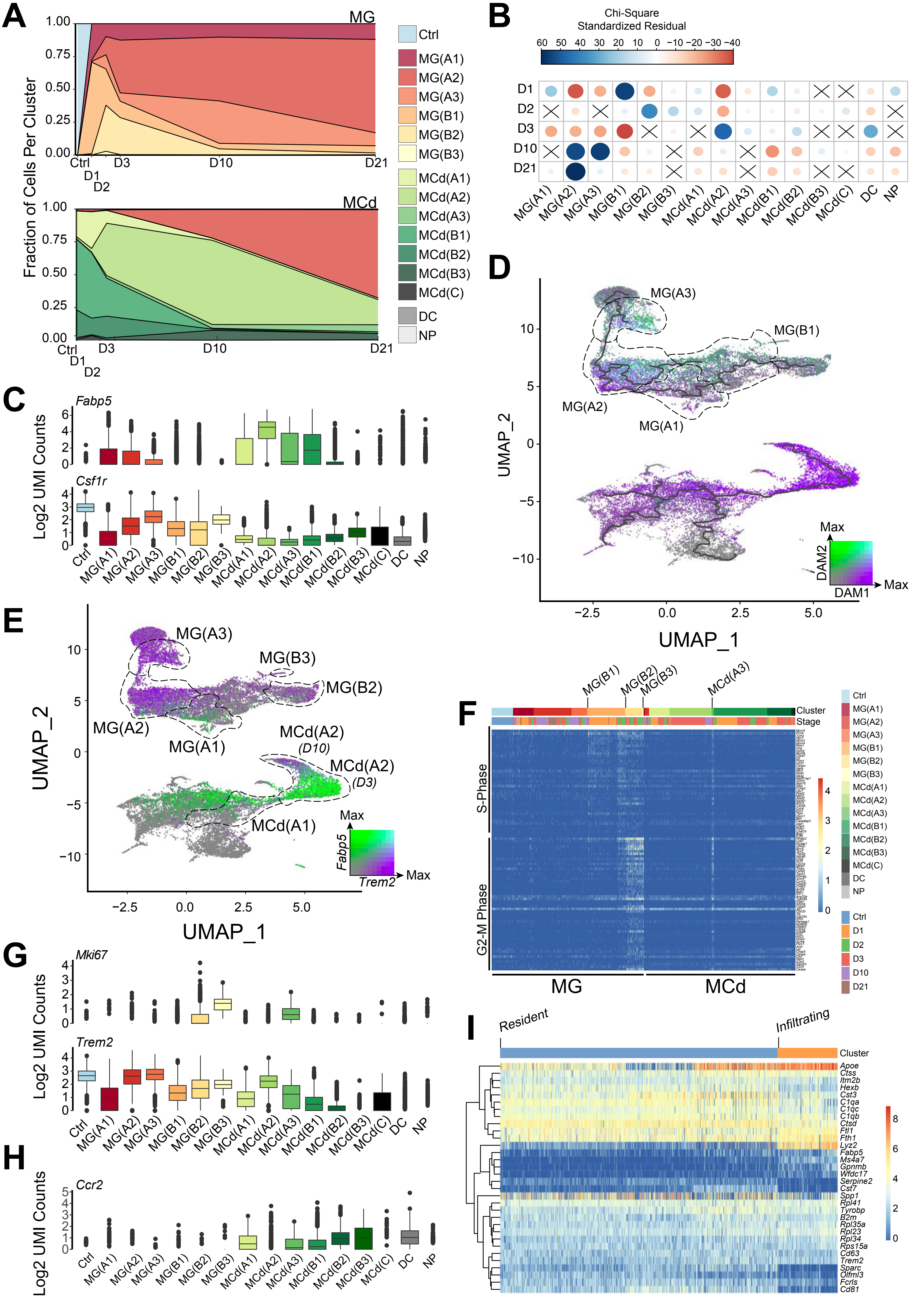

Hamel et al., Extended Data Figure 5

**Extended Data Fig. 5. Clusters represent states of myeloid cells post-SCI. A,** Breakdown of the MG and MCd clusters by the fraction of cells from each time point. **B,** Correlation plot of the Pearson's chi-squared residuals. The size of each circle is proportional to the absolute value of the standardized chi-square residual. Blue signifies a positive standardized chi-square residual, while red signifies the opposite. "X" represents a non-significant contribution. **C,** Boxplots of the $log_2$-transformed counts of *Fabp5* (top) and *Csf1r* (bottom) by cluster. **D,** UMAP coloured by the scaled average expression of DAM1 genes (purple) or DAM2 genes (green) per cell. The progression from DAM1-like to DAM2-like follows the trajectory from MG(A1) to MG(A3). Dotted lines depict relevant clusters. The trajectory is represented by the solid black line. **E,** UMAP coloured by the scaled $log_2$-transformed counts of *Fabp5* (green) or *Trem2* (purple). Dotted lines depict relevant clusters**. F,** Heatmap of the $log_2$-transformed counts from S-phase (top) and G2-M phase (bottom) genes. **G,** Boxplots of $log_2$-transformed counts of *Mki67* (top) and *Trem2* (bottom) by cluster. **H,** Boxplot of the $log_2$-transformed counts of *Ccr2* by cluster. **I,** Heatmap $log_2$-transformed counts for the top 20 differentially expressed genes for MG(A2) plus *Serpine2* and *Gpnmb*. Cells (columns) are labelled by their fate mapping status as in methods

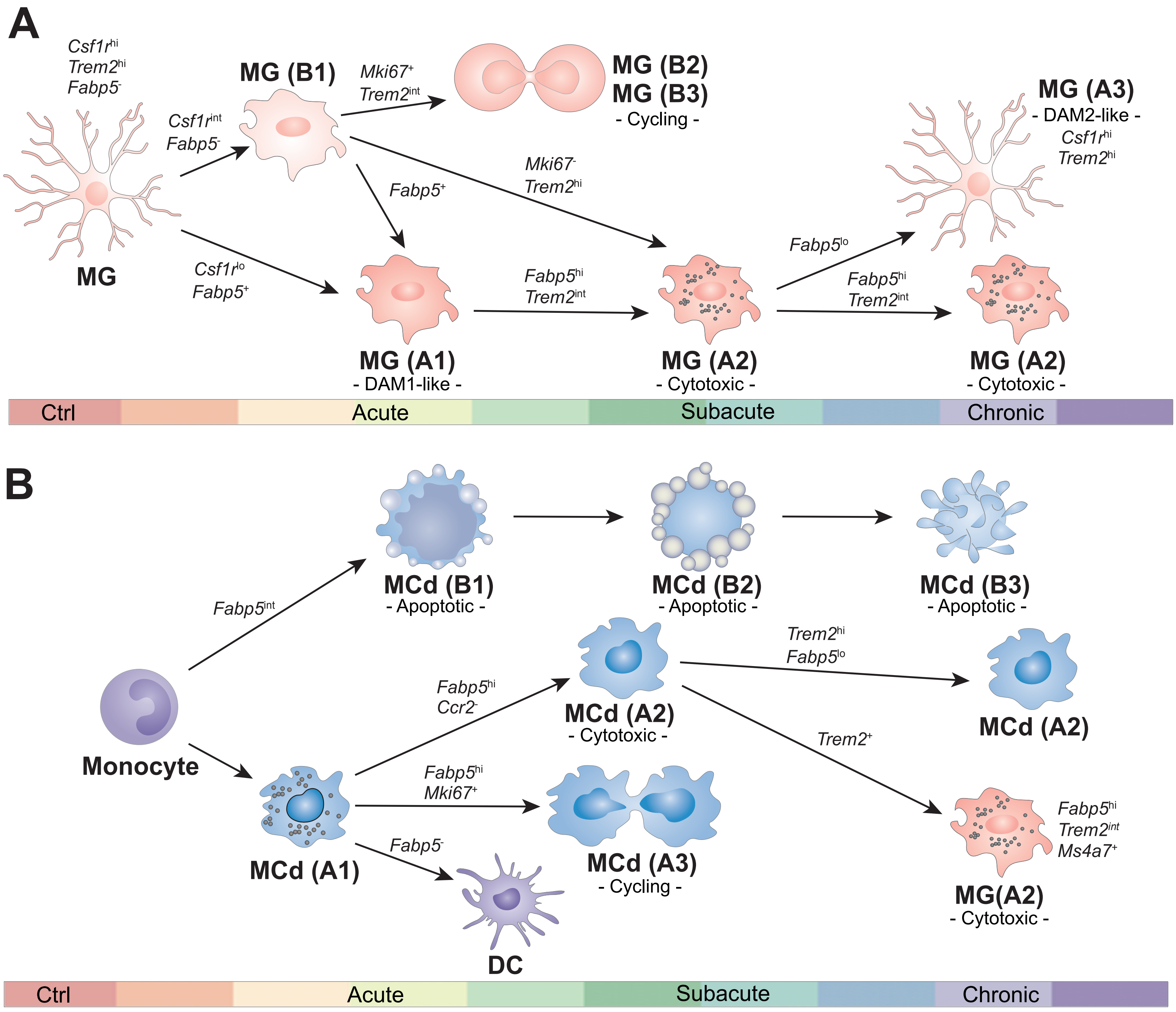

Hamel et al. Extended Data Figure 6

**Extended Data Fig. 6. Proposed map of the myeloid cell trajectories after SCI.**

**A,** MG states over time as in **Fig. 3**. **B,** MCd states over time in **Fig. 3**.

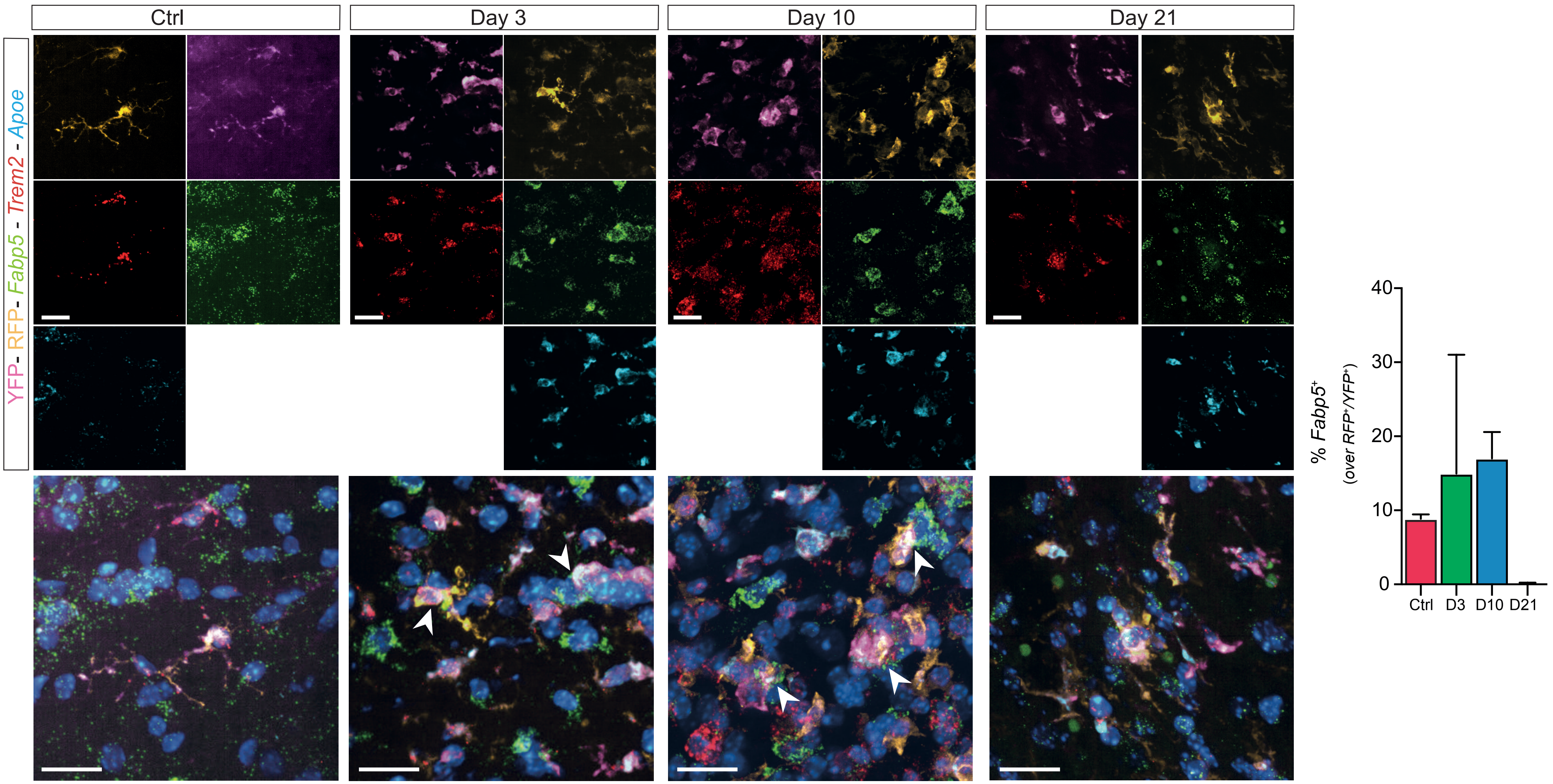

Hamel et al., Extended Data Figure 7

**Extended Data Fig. 7. smFISH validation of data in Fig. 3**. Expression of *Fabp5*, *Trem2* and *Apoe* from Ctrl, D3, D10 and D21. Nuclei in merged panels were stained with DAPI. Arrowheads indicate *Fabp5*$^+$/RFP$^+$/YFP$^+$ cells. Scale bars: 20 μm. The bar graph shows the quantification of *Fabp5*$^+$ cells out of all RFP$^+$/YFP$^+$ CNS resident MG. Data are mean % (± SD) from n= 2 mice per time point.

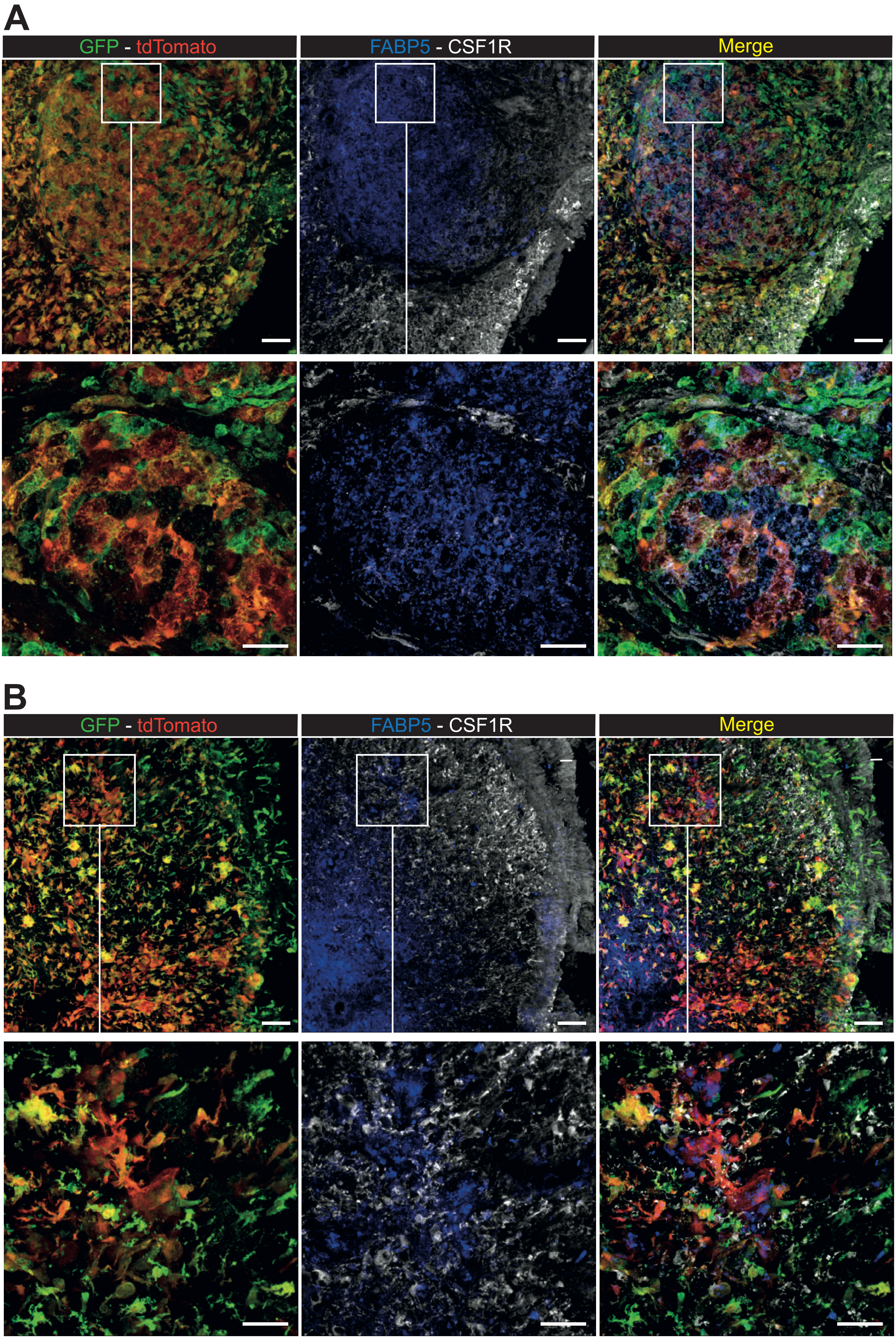

Hamel et al., Extended Data Figure 8

**Extended Data Fig. 8. Confocal immunofluorescence microscopy validation of data in Fig. 3.** D21 expression of FABP5 and CSF1R at the lesion epicentre (**A**) and in the perilesional cord (**B**). Nuclei were stained with DAPI. Scale bars: Top panel 60 μm; Bottom panel 20 μm.

### Summary

| Number of cells in the dataset after quality control | | | | | |
|---|---|---|---|---|---|
| **Total = 30958** | | | | | |
| **Per timepoint** | | | | | |
| **HC** | **1** | **2** | **3** | **10** | **21** |
| 1887 | 8017 | 3858 | 12136 | 3784 | 1276 |
| **Per mouse strain** | | | | | |
| **Cx3cr1CreERT2:R26tdTomato** | | | **Cx3cr1$^{CreERT2}$** | | |
| 17216 | | | 13742 | | |
| **Per sex** | | | | | |
| **Female** | | | **Male** | | |
| 10319 | | | 20639 | | |
| **Stage by Fluorphore** | | | | | |
| | | **Cx3cr1** | **RFP+/YFP+** | **RFP-/YFP+** | |
| | **HC** | 1429 | 458 | 0 | |
| | **1** | 3420 | 0 | 4597 | |
| | **2** | 1354 | 1711 | 793 | |
| | **3** | 4200 | 1219 | 6717 | |
| | **10** | 2218 | 775 | 791 | |
| | **21** | 1121 | 0 | 155 | |
| **Stage by Sex** | | | | | |
| | | **Female** | **Male** | | |
| | **HC** | 1429 | 458 | | |
| | **1** | 1827 | 6190 | | |
| | **2** | 600 | 3258 | | |
| | **3** | 3959 | 8177 | | |
| | **10** | 1383 | 2401 | | |
| | **21** | 1121 | 155 | | |

### Sample Metrics

| SampleID | Type | Sex | Stage | Cell_Count | Median_Genes | Mean_Counts | Mean_Normalized_Counts |
|---|---|---|---|---|---|---|---|
| **SIGAE5** | CreYFP | F | 1 | 1827 | 2726 | 10855.86 | 3.96 |
| **SIGAB11** | CreYFP | M | 1 | 2770 | 2235 | 6866.26 | 3.74 |
| **SIGAF9** | Cx3 | M | 1 | 1239 | 1270 | 3953.37 | 3.51 |
| **SIGAG9.D1** | Cx3 | M | 1 | 804 | 1357 | 4532.20 | 3.57 |
| **SIGAH11** | Cx3 | M | 1 | 1377 | 1078 | 3429.86 | 3.46 |
| **SIGAA1** | CreRFP | M | 2 | 1711 | 1517 | 5566.68 | 3.63 |
| **SIGAF5** | CreYFP | M | 2 | 793 | 2732 | 9758.88 | 3.89 |
| **SIGAF11** | Cx3 | F | 2 | 600 | 1288.5 | 4052.45 | 3.50 |
| **SIGAE11** | Cx3 | M | 2 | 754 | 2864 | 10652.51 | 3.95 |
| **SIGAC11** | CreRFP | F | 3 | 159 | 1403 | 6001.10 | 3.61 |
| **SIGAA7** | CreRFP | M | 3 | 1060 | 877 | 3292.24 | 3.37 |
| **SIGAD11** | CreYFP | F | 3 | 3800 | 2513 | 8541.17 | 3.83 |
| **SIGAB7** | CreYFP | M | 3 | 2917 | 3146 | 11383.07 | 4.01 |
| **SIGAD3** | Cx3 | M | 3 | 253 | 797 | 1923.51 | 3.25 |
| **SIGAE3** | Cx3 | M | 3 | 319 | 805 | 1866.88 | 3.24 |
| **SIGAG9.D3** | Cx3 | M | 3 | 3628 | 1904.5 | 6620.07 | 3.75 |
| **SIGAE3.D10** | CreRFP | F | 10 | 775 | 1266 | 4300.77 | 3.54 |
| **SIGAF3.D10** | CreYFP | F | 10 | 608 | 2039 | 7517.07 | 3.81 |
| **SIGAB10** | CreYFP | M | 10 | 183 | 1097 | 5953.53 | 3.62 |
| **SIGAE10** | Cx3 | M | 10 | 1213 | 1010 | 4635.95 | 3.51 |
| **SIGAF10** | Cx3 | M | 10 | 1005 | 1152 | 5371.64 | 3.57 |
| **SIGAG4** | CreYFP | M | 21 | 155 | 1085 | 7193.73 | 3.64 |
| **SIGAG11.D21** | Cx3 | F | 21 | 1121 | 1101 | 4431.79 | 3.57 |
| **SIGAH2** | CreRFP | M | HC | 458 | 1847.5 | 4047.54 | 3.58 |
| **SIGAD10** | Cx3 | F | HC | 1429 | 1806 | 3917.09 | 3.58 |

**Extended Data Table 1.** Per cell and per sample metrics. Entire dataset after quality control.
